# Identification of distinct functions of GLIS3 in β-cell generation critical to prevention of neonatal diabetes

**DOI:** 10.64898/2026.01.06.697979

**Authors:** David W. Scoville, Sara A. Grimm, Xin Xu, Benedict Anchang, Anton M. Jetten

## Abstract

GLIS3 plays a critical role in pancreatic β cells and diabetes, being one of only a few genes implicated in Type 1, Type 2, and Gestational diabetes. In addition to β cells, GLIS3 is expressed embryonically in bipotent and proendocrine progenitor cells, suggesting a role in endocrine lineage development in mice. We hypothesize that GLIS3 is playing important roles in endocrine development as well as during β cell differentiation. We utilized single cell RNA-sequencing at three different timepoints (embryonic day (e)13.5, e15.5, and e18.5) and single nucleus (sn)ATAC-sequencing at e15.5 in both wild type (WT) and *Glis3* Knockout (*Glis3*KO) mouse embryos. This analysis revealed that the numbers of bipotent, proendocrine progenitor, and all subsequent endocrine cells were proportionately reduced in *Glis3*KO mice. Additionally, loss of GLIS3 function generates a unique subpopulation of cells that fail to upregulate *Ins2* transcription and downregulate expression of several ribosomal and oxidative phosphorylation genes normally repressed during preβ to β cell differentiation. Our study therefore shows that GLIS3 regulates two distinct stages: the embryonic generation/differentiation of bipotent cells and the differentiation of preβ to β cells. Dysregulation of these two stages provides a causal mechanism for the development of neonatal diabetes in GLIS3-deficiency.

## Introduction

The Krüppel-like zinc finger transcription factor, GLI-similar 3 (GLIS3), plays a critical role in the regulation of several biological processes and in the development of various pathologies in humans, including diabetes^1-8^. Within pancreatic β cells, GLIS3 functions as an activator and repressor of gene expression and regulates several critical β cell and diabetes related genes, including the insulin gene, *Ins2*^1,2,9,10^. *GLIS3* is one of a few genes implicated in all main types of diabetes^4-6^, with the loss of GLIS3 function in mice and humans causing neonatal diabetes, and single nucleotide polymorphisms in *GLIS3* associated with an increased risk for Type 1, Type 2, and gestational diabetes in humans^4-6,8,11-16^. The development of neonatal diabetes has been attributed to a decrease in the number of pancreatic β cells and suppression of insulin production^2,5,6,17,18^. Although many insights have been obtained into the transcriptional regulation of insulin gene expression by GLIS3^18-20^, the role of GLIS3 in the regulation of the endocrine lineage and β cell generation during embryonic development remains poorly understood.

Development of the pancreatic endocrine lineage is a multi-stage process^21,22^. During mouse pancreas development, GLIS3 protein is first detectable in SOX9^+^NKX6.1^+^ bipotent progenitor (BiP) cells starting at embryonic day 13.5 (e13.5)^23^. It remains expressed when these cells differentiate into the ductal and proendocrine lineages and continues to be expressed as proendocrine progenitors differentiate into β and pancreatic polypeptide (PP) cells, but becomes suppressed in α, δ, and ε cells. This developmental expression profile suggested that, in addition to its role in β cells, GLIS3 might have additional roles during embryonic pancreatic development. This is supported by studies showing a reduction in the number neurogenin 3-positive (NGN3^+^) proendocrine progenitors in *Glis3*-knockout mice (*Glis3*KO)^24,25^. In humans, a similar phenotype is observed in human embryonic stem cells differentiated to β-cells, with cells lacking functional GLIS3 producing fewer β-cells^26^. However, whether loss of GLIS3 function affects the generation of BiPs, their differentiation into proendocrine progenitors, and to what extent it regulates subsequent stages in the endocrine lineage has yet to be determined. Human stem cell models have been insufficient to address these questions.

To obtain insights into the role of GLIS3 in the regulation of the different stages of prenatal pancreatic endocrine development and how it relates to the causal mechanisms of neonatal diabetes in GLIS3-deficiency, we analyzed gene expression in different pancreatic subpopulations by single cell RNA-sequencing (scRNA-Seq) analysis. We examined cells from 3 different timepoints (e13.5, e15.5, and e18.5), which allowed us to model pancreas development and examine GLIS3 function at each stage. To examine whether GLIS3 affected chromatin accessibility of target genes single nucleus ATAC-seq (snATAC-seq) was performed at a single timepoint (e15.5). Our study demonstrates that loss of GLIS3 function regulates two distinct stages during endocrine development that together provide the causal mechanism of neonatal diabetes in GLIS3 deficiency.

## Materials and Methods

### Mouse Strains

The *Glis3*KO strain, B6.129S-Glis3<TM1AMJ>, in which exon 5 of *Glis3* containing the third and fourth zinc finger motifs of the DNA binding domain was deleted, was described previously^2,27^. Mice were maintained on a C57BL/6 background and routinely fed an NIH-31 diet (Harlan). For embryonic age, the morning of vaginal plug discovery was considered e0.5. Sex was established by genotyping for the Y chromosome. All animal studies followed the guidelines outlined in the NIH Guide for the Care and Use of Laboratory Animals, and the protocols were approved by the Institutional Animal Care and Use Committee at the National Institute of Environmental Health Sciences (NIEHS).

### ScRNA-seq and snATAC-seq

Embryonic pancreas was dissected at e13.5, e15.5, and e18.5. Pancreata were dissociated into single cells by digestion with TrypLE Express Enzyme (Gibco, Waltham, MA) for 30-60 minutes at 37°C, with pipette trituration every 15 minutes. Trypsinization was stopped by adding an equal volume of cold fetal bovine serum (FBS, GeminiBio, West Sacramento, CA), Cells were subsequently resuspended in 90% FBS/10% DMSO and stored at -80°C for up to 3 months while sex and genotype were established. Cells were then thawed, washed with media containing 10% FBS in DMEM high glucose with penicillin/streptomycin (Gibco, Waltham, MA), passed through a 35 μm strainer, and prepared for single cell sequencing or single nucleotide sequencing following the 10X Genomics protocol (10X Genomics, Pleasanton, CA). Male embryos were used for sequencing as they were the first to be obtained in sufficient numbers. For scRNA-seq at e13.5, 3-4 embryonic pancreata were pooled for each WT and KO sample. For scRNA-seq at e15.5 and e18.5, one embryonic pancreas was used for each WT and KO sample. For snATAC-seq at e15.5, 1-2 pancreata were pooled and used per sample. Samples were processed on a Chromium X (10X Genomics, Pleasanton, CA), following manufacturers protocol. The scRNA-seq and snATAC-seq libraries were sequenced on an Illumina NovaSeq 6000 by the NIEHS Epigenomics and DNA Sequencing Core, with ∼100,000 reads obtained per cell.

### Data Analysis

Counts per gene for the scRNA-seq samples were collected by 10x Genomics Cell Ranger count (v6.0.0)^28^ using the “--include-introns” option, with gene models defined by GENCODE M17. Seurat (v4.1.0)^29^ was used to perform an initial filtering of low-quality cells with thresholds applied at 15% mitochondrial gene content, 0.5% hemoglobin gene content, and a minimum of 1000 detected genes per cell. Putative doublet cells were removed via scDblFinder (v.1.1.8)^30^. Ambient RNA decontamination was deemed unnecessary for this dataset, as the predicted ambient RNA levels by SoupX (v1.5.2)^31^ were low (2%-6% per sample). Seurat (v4.1.0) was used to carry out normalization and sample-based integration for all cells passing the previously listed filters. Broad cell type annotations were assigned to clusters based on expression of known marker genes. Cells identified as endocrine cells were extracted from the full dataset and re-integrated based on age due to low cell counts in several samples. Endocrine subtype annotations were assigned to endocrine cell subclusters (at resolution=0.4) based on the expression of known marker genes. Differentially expressed genes were identified via Seurat’s FindMarkers function with logfc.threshold set to 0. Post-processed nCount and nFeature data for each sample were similar between samples (Supplemental Figure 1A-B).

The raw snATAC-seq samples were pre-processed with 10x Genomics Cell Ranger ATAC count (v2.1.0)^32^ using the mm10 reference pre-built for Cell Ranger ARC (version 2020-A-2.0.0) to identify initial open chromatin regions per sample. These peak calls were collapsed to get a single combined peak set for the full dataset, ignoring any peaks that were <20 bp or >10,000 bp or located on a non-canonical chromosome. Counts per peak were collected over the combined peak set via Signac (v1.6.0)^33^. Prior to integration by sample of the snATAC-seq data in Signac, the nuclei were filtered with the following requirements: >25% of reads in peaks, <5% of reads in blacklist regions, TSS enrichment score >2, aggregate reads at peaks >500, and aggregate reads at peaks <50,000. Broad cell type annotations were transferred from the e15.5 scRNA-seq analysis based on cis-co-accessible networks (CCANs) determined with Cicero (v1.3.6)^34^. The predicted endocrine cluster was extracted from the full snATAC-seq dataset, and a new set of open chromatin peak calls was made by MACS2 (v.2.1.1)^35^ using the Signac CallPeaks function with the ‘group.by’ parameter set to “Sample”. Signac was again used to collect counts per peak, this time for only the predicted endocrine nuclei and using the endocrine-specific peak set, followed with sample integration by genotype. Cell type annotations, again based on CCANs from Cicero, suggested that a subset of nuclei in the endocrine cluster likely originated from acinar rather than endocrine cells. After the presumed acinar nuclei were removed, integration and annotation transfer were repeated, this time referencing the endocrine cell subtypes instead of the broad cell types in the e15.5 scRNA-seq analysis. Peaks with differential accessible chromatin signal were identified via Seurat’s FindMarkers function with the following settings: min.pct = 0.05, test.use = “LR”, latent.vars = "peak_region_fragments", and logfc.threshold = 0). ScRNA-Seq and snATAC-Seq analyses were submitted to the Gene Expression Omnibus under access numbers GSE285200 and GSE285201, respectively.

## Results

### Glis3KO mice generate fewer endocrine cells

To obtain greater insights into the regulation of the endocrine lineage by GLIS3 during pancreas development, pancreata from different embryonic stages were analyzed by scRNA-seq. As GLIS3 protein expression is first detected at e13.5 in the mouse pancreas^23^, we collected samples at e13.5, e15.5, and e18.5 to allow for enrichment of cells at both early and late stages in endocrine lineage development (N=3 WT and KO, at each timepoint). After sequencing, all embryonic cell data were combined to produce enough cells in each subpopulation at each developmental stage for analysis. Cell type identities of broad clusters (Figure 1A) were assigned based on the expression of known marker genes (Figure 1B). Cell numbers in each cluster were then quantified and normalized to the number of sorted cells passing quality filters (Figure 1C). While no significant differences were observed in cell number of most cell populations between WT and KO mice, including acinar and Stellate cells, pancreata from KO mice contained significantly fewer total endocrine cells than WT mice (1.6% vs 5.7%, Figure 1C). Separate cell counts for e13.5, e15.5 and e18.5 revealed that the difference in total endocrine cell number was first apparent at e13.5 but reached statistical significance at e15.5 (Supplemental Figure 2A-D) suggesting that GLIS3 regulates an early stage of embryonic endocrine lineage development. Although few acinar cells were detected at e18.5 in both WT and *Glis3*KO samples, likely due to technical issues of over-trypsinization, this did not change the conclusion that the number of endocrine cells was decreased at all 3 times point analyzed (Supplemental Figure 2D). That GLIS3 did not affect acinar cells is consistent with previous studies showing that loss-of-GLIS3 function has no significant effects on pancreatic acini development and acinar gene expression^2,5,23,36^.

**Figure 1.**
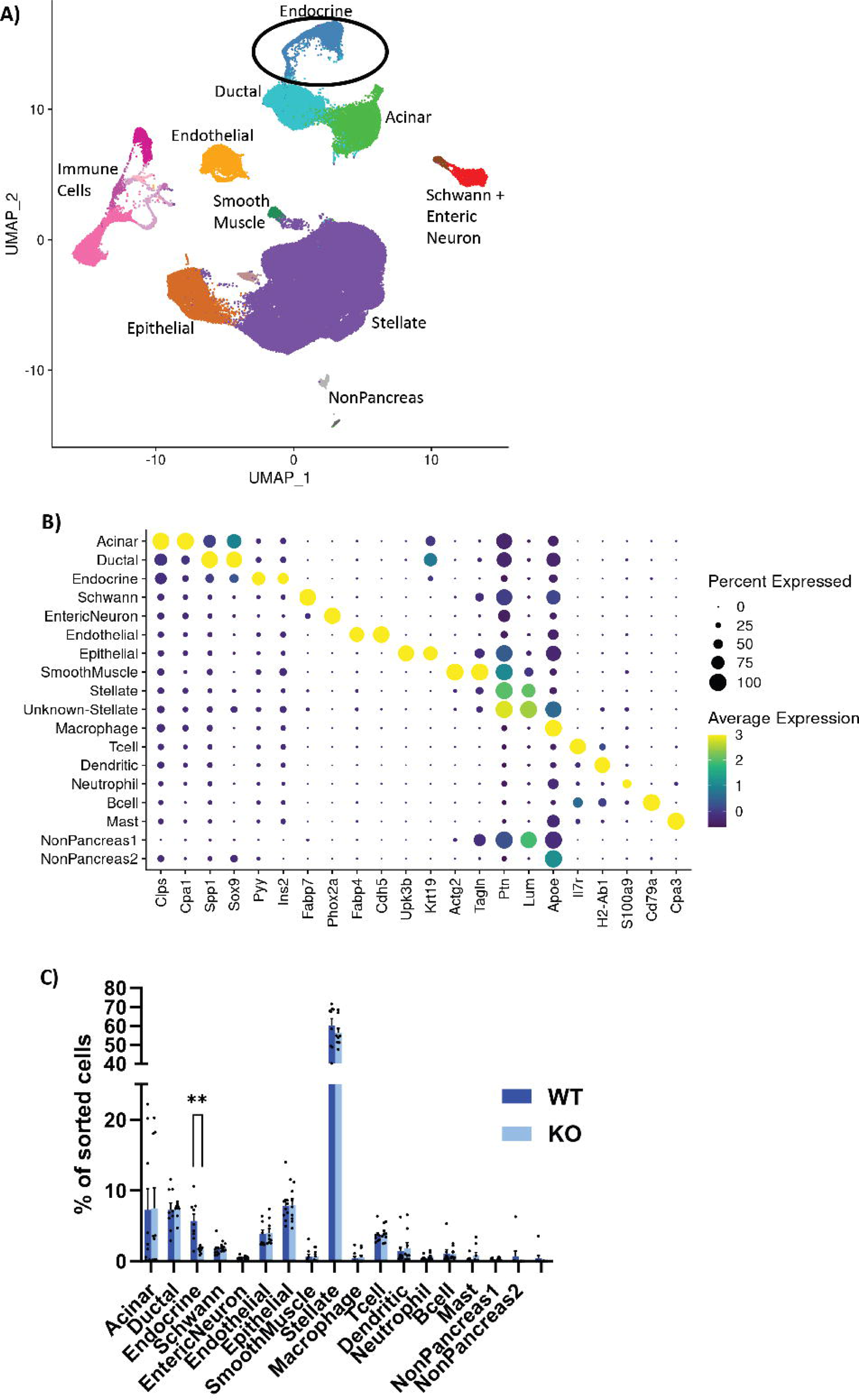
ScRNA-seq reveals reduction in endocrine cell population in embryonic pancreas. (**A)** UMAP clusters of combined pancreatic cells from e13.5, e15.5, and e18.5. (**B)** Dotplot of marker genes for specific clusters, with color indicating the level of expression and size indicating the percentage of cells expressing that gene within the cluster. (**C)** Quantification of cells in each subcluster, as a percentage of the total cells sorted per sample, separated by genotype. N=9 for WT and KO, error bars indicate SEM, ** indicates p<0.01.

### GLIS3 regulates two distinct stages of pancreatic endocrine development

Previous studies indicated that the reduction in the number of pancreatic β cells and decreased insulin production/secretion are major factors in the development of neonatal diabetes in mice and humans with GLIS3 deficiency^2,5,17,23,37^. To obtain greater insights into the functions of GLIS3 in pancreas development, we focused our scRNA-Seq analysis on the endocrine lineage, as well as BiP cells (Figure 2A). BiP cells clustered with endocrine cells in our analysis and were identified by their expression of *Nkx6.1* and *Sox9* (Figure 2B and C). Pro-endocrine cells were identified by their expression of *Neurog3* (Figure 2D), and late pro-endocrine FEV+ cells by their expression of *Fev* (Figure 2E). Beta and Delta cells were identified by their increased expression of *Ins2* and *Sst*, respectively (Figure 2F and G). Alpha and Epsilon cells did not subcluster separately, although the pattern of expression of *Gcg* and *Ghrl* appeared mostly distinct (Figure 2H and I). The Alpha2 subcluster came primarily from e13.5 samples, indicating that these cells may represent a first wave of differentiation, which primarily produces Alpha cells (Figure 2J) ^38,39^. As expected, most of the cells in the earliest stages of differentiation (BiP Cells and Pro-Endocrine cells) came from the e13.5 and e15.5 samples, whereas most of the later stages of hormone-positive cells came from the e18.5 samples (Figure 2J and Supplemental Figure 3).

**Figure 2.**
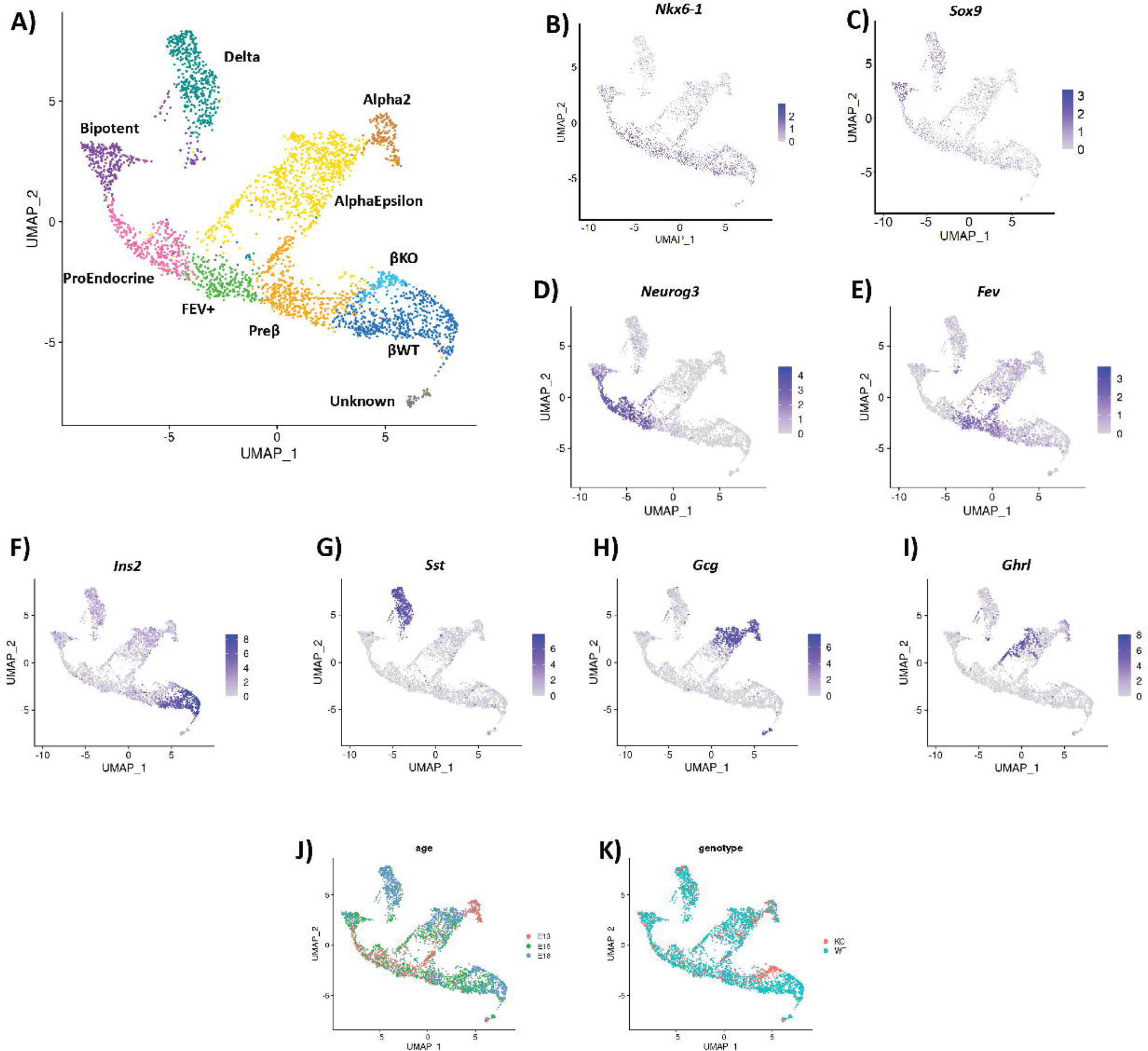
WT and *Glis3*KO endocrine cells largely overlapped, except for β cells. (**A**) UMAP of the endocrine cell subcluster from Figure 1A. (**B-E)** Cell clusters were identified based on their expression of *Nkx6.1*, *Sox9*, *Neurog3*, *Fev*. (**F-I)** Cell clusters were identified based on their expression of *Ins2*, *Sst*, *Gcg*, and *Ghrl*. (**J**) Cells are labeled by age, with more of the e13.5 cells present earlier in development and more e18.5 cells in the hormone-positive clusters. (**K)** Genotype information showed that the WT and KO cells largely overlapped, except for the β cell subclusters.

Comparison of WT and *Glis3*KO cells in the scRNA-Seq analysis revealed that although the total number of endocrine lineage-related cells (including BiP cells) was significantly reduced (70%) in *Glis3*KO mice (Figure 3A), the percentages of each distinct subpopulation were similar between WT and KO cells (Figure 3B). This supports the hypothesis that the decrease in BiPs in *Glis3*KO pancreas consequently affects the generation of all subsequent stages of the endocrine lineage proportionally and that loss of GLIS3 function does not alter lineage determination of proendocrine cells along the α, β-like, δ, and ε endocrine lineages. Additional support for this hypothesis comes from the analysis of the BiP, proendocrine, and endocrine lineages of WT and *Glis3*KO pancreata, which revealed largely overlapping subclusters, except for the β cell subcluster, indicating a critical function for GLIS3 in the regulation of this stage of the β cell lineage (Figure 2A and K).

**Figure 3.**
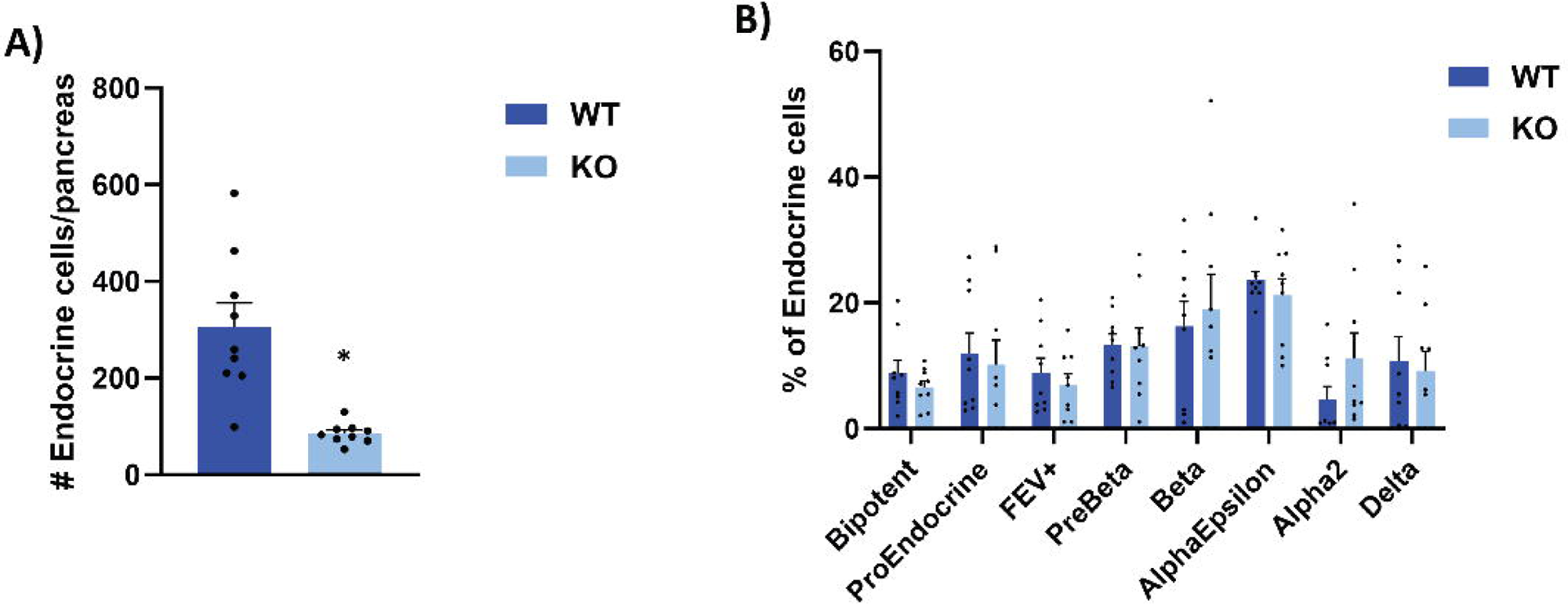
*Glis3*KO mice have proportionally similar numbers of endocrine subtypes. (**A**) Total number of endocrine cells/pancreas in WT and KO embryos. (**B**) The percentage of endocrine cells that were assigned to specific subclusters for each genotype is shown. N=9 for WT and KO, error bars indicate SEM. * indicates p < 0.05.

### Glis3KO pancreas contains a unique β cell subpopulation

The distinct WT and *Glis3*KO β cell subclusters identified here were referred to as βWT and βKO, although a few KO cells were present in the βWT cluster and vice versa. Like βWT cells, βKO cells were primarily observed at e15.5 and e18.5. Comparison of differentially expressed genes between WT and KO within the distinct endocrine subpopulations revealed few differences, except for the βWT and βKO cell populations, consistent with the UMAP data (Table 1, Supplemental Tables 1-10, and Figure 2K). One of the major differences between βWT and βKO was the expression of *Ins2*, with *Ins2^+^* β cells almost entirely restricted to βWT cells (Figure 2F). This decrease in *Ins2* expression in βKO cells is consistent with previous studies showing that GLIS3 plays a critical role in the regulation of *Ins2* transcription^18-20^. Although *Ins2* expression differed between these clusters, they shared expression of many other β cell markers, including *Slc2a2, Iapp,* and *G6pc2* (Supplemental Figure 4). Moreover, in UMAP the βKO cluster lies near the Preβ and βWT cluster, indicating that they are closely related and supporting our conclusion that the βKO cluster represents a distinct β-like cell type.

**Table 1.** KO vs WT genes differentially regulated in different endocrine.

|  | **<u># Gene Differences</u>** | **<u>Upregulated</u>** | **<u>Downregulated</u>** |
| --- | --- | --- | --- |
| **Bipotent** | **11** | **2** | **9** |
| **ProEndocrine** | **9** | **5** | **4** |
| **FEV+** | **2** | **0** | **2** |
| **Preβ** | **18** | **8** | **10** |
| **βKO vs βWT** | **645** | **623** | **22** |
| **Alpha2** | **0** | **0** | **0** |
| **AlphaEpsilon** | **10** | **8** | **2** |
| **Delta** | **36** | **29** | **7** |
| **subpopulations.** |  |  |  |

Volcano plots of differentially expressed genes illustrate that many of the genes downregulated during the Preβ to βWT transition remained expressed at a higher level in βKO cells (Figure 4A, B). ScRNA-Seq analysis identified 2248 changes in gene expression between Preβ and βWT, most of which (∼90%) consisted of genes downregulated in βWT cells, with KEGG analysis linking many of these genes to the ribosome and oxidative phosphorylation pathways (Supplemental Table 11). Comparison of the gene expression profiles between βKO and βWT populations showed that most differentially expressed genes were found to be expressed at a higher level (623 genes) in the βKO subcluster compared to βWT cells, while only 22 genes (including *Ins2*) were suppressed (Figure 4A and Table 1). These dual effects on gene expression are consistent with reports that GLIS3 can function as an activator and repressor of gene transcription^2,17,19,24,40,41^. Interestingly, pathway analysis of genes expressed higher in βKO than in βWT revealed that many were associated with the ribosome and oxidative phosphorylation pathways (Supplemental Table 12). About 30% (189 genes) of the genes with higher expression in βKO cells were downregulated in normal development (Preβ vs βWT)(Figure 4C), and again many of these overlapping genes were linked to ribosome and oxidative phosphorylation pathways (Supplemental Table 13) suggesting that GLIS3 likely regulates the normal development of β cells in large part through suppression of these pathways. Together, these observations indicate that loss of GLIS3 function significantly represses a distinct part of the Preβ to β cell transition and establishes a unique, more immature β cell subpopulation.

**Figure 4.**
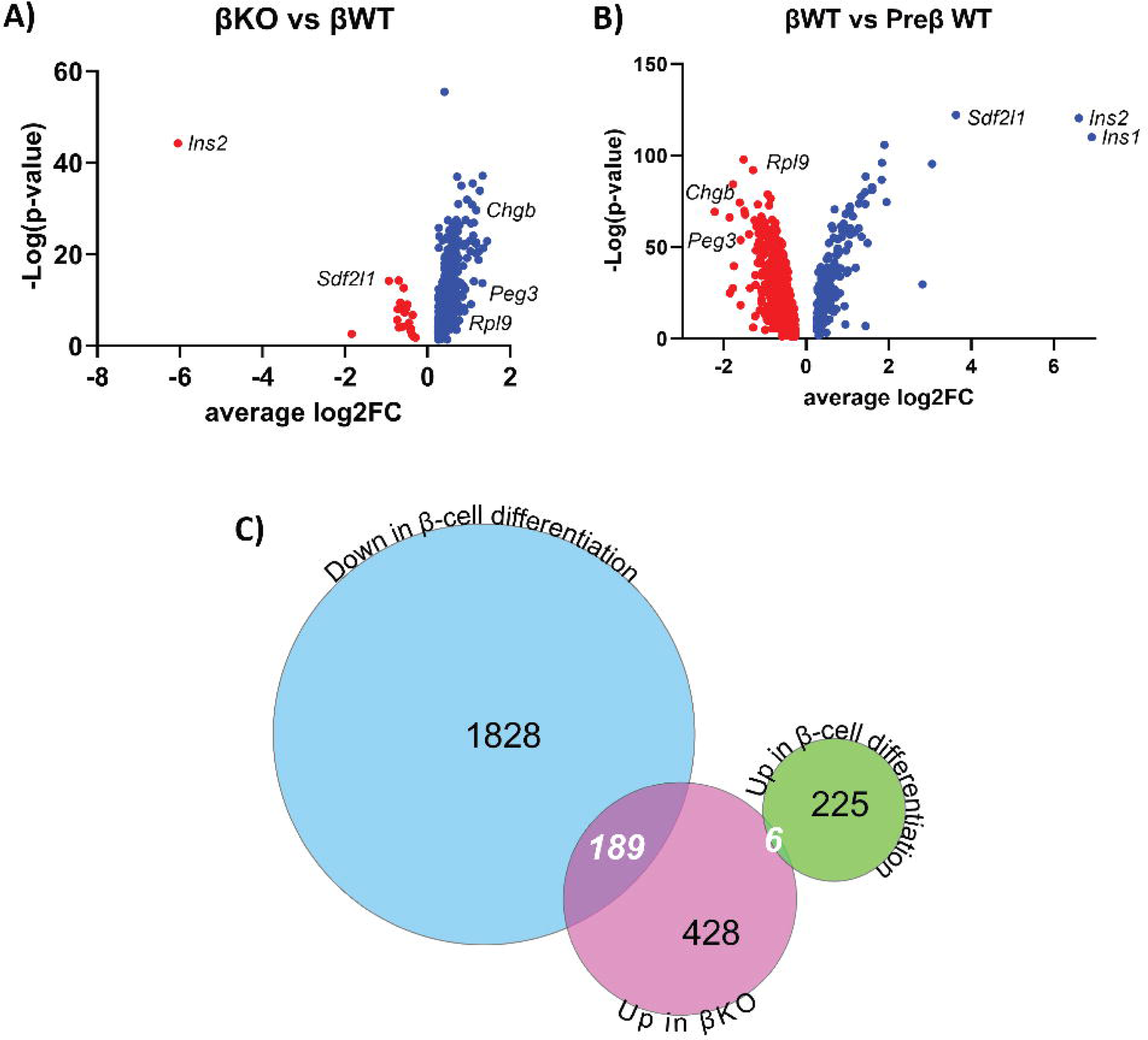
Many genes differentially expressed between WT and KO β cells are developmentally regulated. (**A**) Volcano plots of genes significantly differentially expressed between βKO and βWT subclusters (red, expressed at lower levels; blue, expressed at higher levels). (**B**) Volcano plots of genes significantly differentially expressed between βWT and Preβ WT subclusters. (**C**) Venn diagram comparing genes significantly upregulated in *Glis3*KO β cells (red circle, 623 genes) to genes either downregulated (blue circle, 2017 genes) or upregulated (green circle, 231 genes) during normal development from Preβ to β cells.

In addition to inhibition of the Preβ to β cell transition, several of the genes upregulated in βKO cells did not follow a clear developmental pattern. Some genes (e.g., *Kcnip4, Gatm*) with little known function in pancreas had minimal-to-no expression in the Preβ or βWT cells but were induced in βKO cells (Figure 5A-B). Other genes (e.g., *Rbp4*, *Nnat*, *Iapp*) were upregulated during the Preβ to β transition but were expressed at even higher levels in βKO (Figure 5C-E). The expression of *Chgb*, encoding chromogranin B protein^42^, is normally induced in FEV+ cells, maintained in Preβ cells, and then reduced in β cells, but remained expressed at a higher level in βKO compared βWT cells (Supplemental Figure 5). These differences in gene expression highlight the complexity of the *Glis3*KO β cell phenotype and suggests that GLIS3 plays a critical and diverse role in maintaining β cell identity apart from its role in regulating known pathways (i.e. oxidative phosphorylation) or *Ins2* transcription.

**Figure 5.**
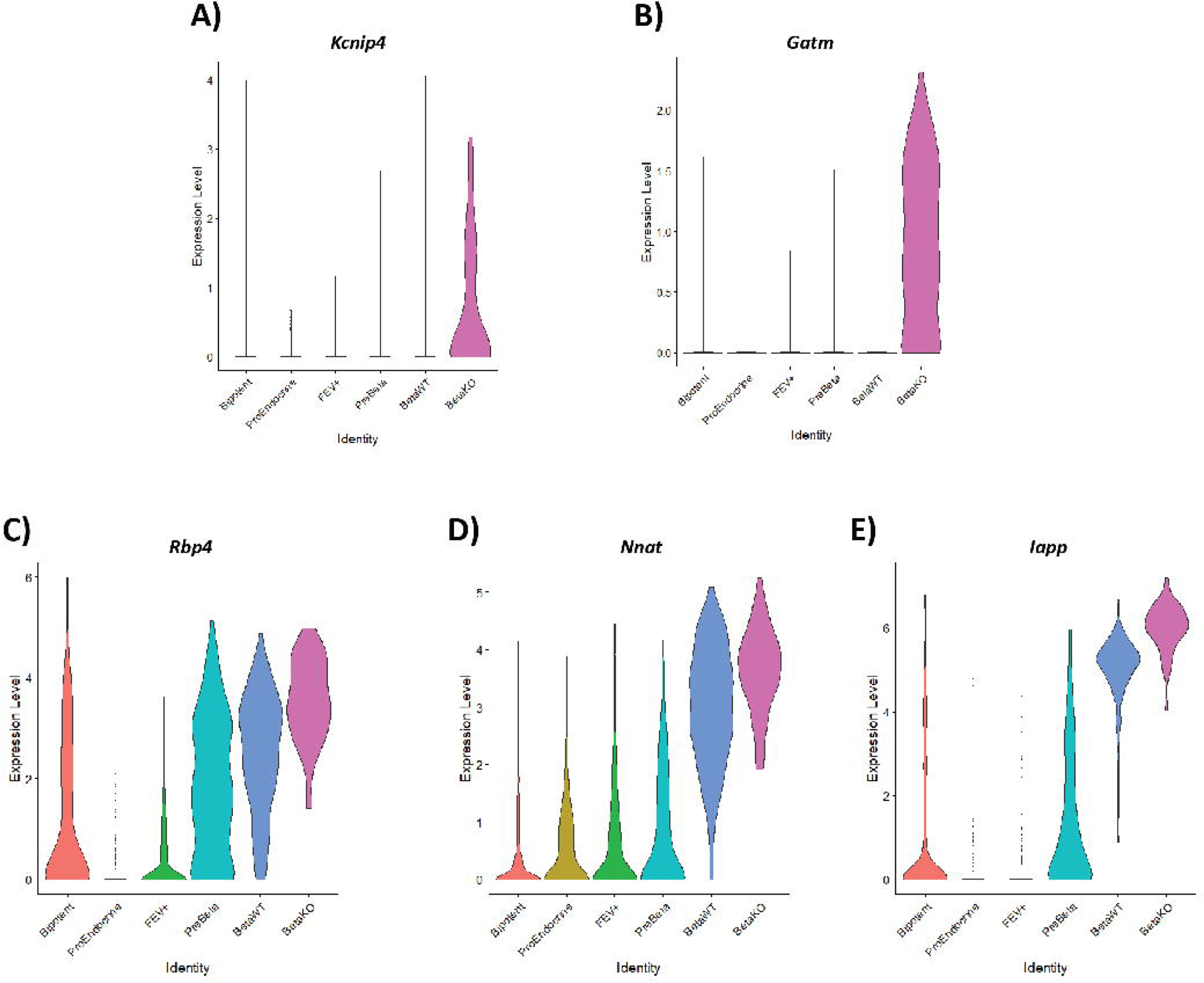
A subset of genes is induced at a higher level in βKO compared to βWT cells during the Preβ to β cell differentiation. (**A-B**) Violin plots showing high induction of genes in βKO that are not in progenitor or βWT cells. (**C-E**) Violin plots showing significantly higher expression of several genes during the Preβ to β transition in βKO compared to βWT cells.

### GLIS3 acts as a cooperating transcription factor in regulating Ins2 transcription

While determining direct binding of GLIS3 to regulatory regions of these genes during early embryonic development is not currently technically feasible, previously published GLIS3 ChIP-seq data from postnatal islets (primarily made up of β cells) indicated that many of these genes, including ribosomal and oxidative phosphorylation genes, are directly bound by GLIS3^19^. The critical role GLIS3 plays in the regulation of *Ins2* transcription is well established^2,3,18,19,24,41^ and as we show in this study GLIS3 regulates the expression of *Ins2* during the Preβ to β cell transition (Figure 2F). To examine whether GLIS3 directly controls chromatin accessibility at the *Ins2* locus or that of other genes regulated by GLIS3, we performed snATAC-seq on pancreata isolated from e15.5 WT and *Glis3*KO embryos. Cell type predictions were made using Cicero, comparing snATAC-seq to e15.5 scRNA-seq data, and the endocrine subcluster identified in snATAC-seq (Figure 6A). Most of the expected endocrine subpopulations were represented in the snATAC-seq data, apart from Delta cells, which were relatively few at e15.5 (Figure 2A and 2J). Interestingly, the WT and KO β cell populations did not differ dramatically (Supplemental Figure 7), as was observed in the scRNA-seq data. Comparison of snATAC-seq signals between βKO cells and βWT cells revealed few, relatively small changes in signal (Supplemental Table 14). Since *Ins2* is the gene most significantly transcriptionally regulated by GLIS3, we focused on the *Ins2* locus. Chromatin accessibility surrounding *Ins2* was mostly closed during early endocrine development but opened after differentiation into βWT cells (Figure 6B). Comparison of the ATAC signal at the *Ins2* locus between βWT and βKO cells showed that the increase in ATAC signal in βKO was smaller compared to that in βWT cells (Figure 6B). In contrast, examination of the ATAC signals at the *Chgb* and *Sdf2l1* loci, genes that were, respectively, highly upregulated or downregulated in βKO cells, showed no noticeable difference in chromatin accessibility at these loci between *Glis3*KO and WT mice at all stages of the embryonic endocrine to β lineage (Figure 6C and D). These data suggest that GLIS3 is not required for opening closed chromatin at these loci and therefore does not act as a pioneer factor but further enhances chromatin accessibility at the *Ins2* locus. This suggests that GLIS3 may act as a cooperating transcription factor, as in a recently proposed model of transcriptional regulation^43^, and in coordination with a pioneer transcription factor increases chromatin accessibility at the *Ins2* promoter allowing dramatic activation of *Ins2* transcription (Figure 6E).

**Figure 6.**
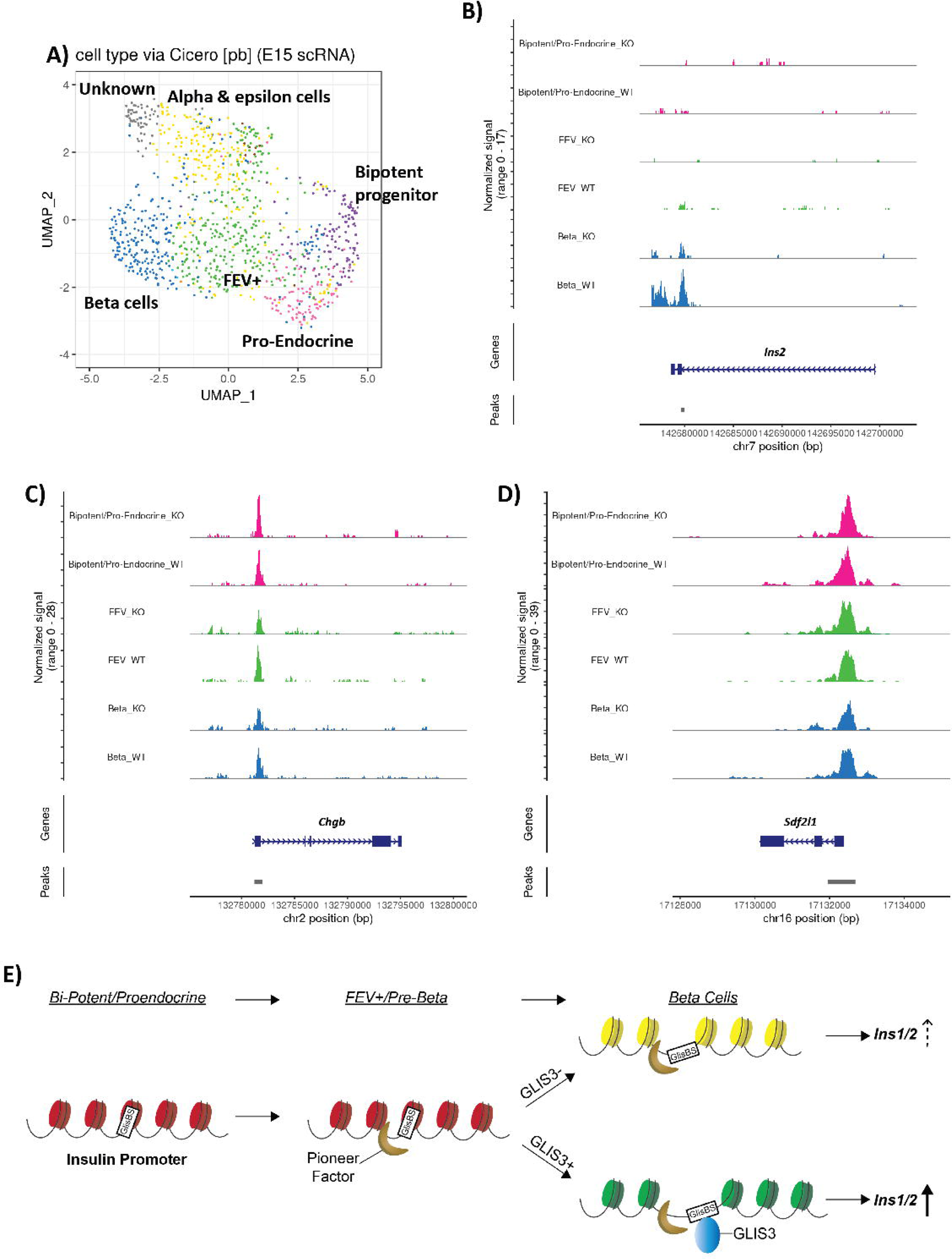
SnATAC-seq at e15.5 suggests GLIS3 does not function as a pioneer factor. **(A**) UMAP showing the endocrine subcluster of snATAC-seq data, with cell identities determined via Cicero. Genomic tracks for ATAC-seq data from the indicated subclusters, with BiP and Pro-endocrine cells being combined to one track, for the (**B**) *Ins2*, (**C**) *Chgb*, and (**D**) *Sdf2l1* genes. (**E**) Model of GLIS3 binding to the insulin promoter and acting as a cooperating transcription factor. GlisBS indicates a GLIS3 binding site.

## Discussion

Previous reports have suggested that a loss of GLIS3 function leads to neonatal diabetes through a combination of a decrease in the number of pancreatic β cells and suppression of insulin gene transcription and production^2,5,6,17,18^. In this study, we provide greater insights into the causal mechanism of neonatal diabetes and show that during embryonic pancreas development, GLIS3 regulates two distinct stages of the endocrine lineage. GLIS3 protein is first detectable in BiP cells^23^, and when these cells differentiate along the ductal and endocrine lineages, GLIS3 remains expressed in the ductal cells and proendocrine progenitors, as well as in β and pancreatic polypeptide (PP) cells, but not in α, δ, and ε cells^23^. This was supported by our scRNA-seq analysis (Supplemental Figure 6A-C). The expression of GLIS3 during embryonic pancreas development suggested that GLIS3 may have a regulatory role in the generation, maintenance and/or differentiation of BiP cells and/or the subsequent endocrine lineages. This is supported by our scRNA-Seq analysis showing that the total number of endocrine cells was 75% lower in *Glis3*KO pancreata than those of WT; however, no significant differences were observed in the relative number of BiP, proendocrine, and the different endocrine populations between WT and *Glis3*KO pancreata (Figure 3A and Figure 3B). These data indicate that loss of GLIS3 function reduces the generation of BiPs and all endocrine cells similarly and supports the concept that the decrease in BiPs in *Glis3*KO pancreas subsequently resulted in a proportionate decrease in all succeeding stages of the endocrine lineage. That BiPs are the first cell type in which GLIS3 is expressed during pancreas development would be consistent with a function of GLIS3 at this stage.

When proendocrine cells differentiate along the different endocrine lineages, GLIS3 remains expressed in β and PP cells and is repressed in α, δ, and ε cells^2,3,24^. This raised the possibility of a role for GLIS3 in endocrine lineage determination. However, our data shows no difference in the relative number or UMAP positioning of the endocrine subpopulations between WT and *Glis3*KO pancreata (Figure 2K and 3B), indicating that the loss of GLIS3 function does not alter the determination of proendocrine cells to differentiate along the different endocrine cell lineages. Neurogenin 3 (encoded by *Neurog3*) has been reported to play a critical role in the differentiation of BiPs into proendocrine progenitors^21,22,44-47^, while other studies provided evidence for a role of GLIS3 in the regulation of *Neurog3* transcription^2,3,24^. These observations led to the conclusion that loss of GLIS3 function inhibits the differentiation of BiPs into proendocrine cells by suppressing *Neurog3* expression, which subsequently is responsible for the reduction in β cell generation. However, our scRNA-seq analysis indicated that the expression of *Neurog3* in proendocrine progenitors was not significantly different between WT and *Glis3*KO mice (Supplemental Table 2), suggesting that *Neurog3* is not regulated by GLIS3. This conclusion is consistent with our ChIP-Seq analysis showing that GLIS3 does not bind to the *Neurog3* locus^19^. Our data indicates that the decreased *Neurog3* expression seen in studies with whole pancreas rather reflects the decrease in proendocrine cells caused by the reduction in BiP cells.

Although the number of endocrine cells is reduced, the number of ductal cells was similar between WT and KO (Figure 1C), which was surprising considering that they are also derived from BiP cells, and thus one would expect them to also be reduced in number. We hypothesize that this is caused by a combination of shunting of BiPs to the ductal lineage and/or an increase in proliferation in early ductal cells in the absence of functional GLIS3. The latter is consistent with our previous observations in *Glis3*KO and pancreas specific knockout of *Glis3*, where we observed cystic pancreatic ducts^2,19^, indicating an increase in ductal cell proliferation. Alternatively, it is possible that the cells traditionally referred to as BiPs are clustering within the ductal cell cluster, and what we have labeled BiPs here represent a transient stage in between traditional BiP and pro-endocrine stages. However, a lack of known individual marker genes for BiPs and limitations in BiP cell number in our analysis precludes us from further distinguishing these subpopulations. Regardless, the data presented here support a critical and previously unappreciated role for GLIS3 during the initial stages of BiP differentiation to pro-endocrine cells that warrants future study.

While loss of GLIS3 function does not alter the determination of proendocrine cells along the different endocrine lineages, it significantly changes the Preβ to β cell trajectory and produces a unique β cell-like population during embryonic development. Thus, in addition to acting at the BiP stage, GLIS3 is critical for the Preβ to β cell transition. Transcriptome analysis identified 645 genes that were differentially expressed between the βKO and βWT populations (Table 1). A vast majority (623) of these genes were expressed at a higher level in βKO cells, consistent with our previous studies in islets from 4-week-old mice with a pancreas-specific deletion of *Glis3* showing roughly twice as many upregulated as downregulated genes^19^. Interestingly, only ∼25% of these upregulated genes continued to be have higher expression in *Glis3* KO postnatally^19^. Our study shows that these defects observed in postnatal *Glis3*KO β cells are the result of changes made during embryonic development when Preβ cells transition into β cells (Figure 2A). In WT mice, nearly 10 times as many genes are repressed as activated during the Preβ to β cell transition (Figure 4B), while in *Glis3*KO mice the downregulation of many of these genes is suppressed (Figure 4C). Interestingly, many of these genes are ribosomal subunits or related to oxidative phosphorylation. Specific ribosomal subunits have been reported to be associated with the increased translation of specific mRNAs, such as the translation of a subset of *Hox* genes by RPL38^48^, and changes in ribosomal subunit expression have been linked to several cellular functions^49^ and disease states^50^. However, roles for specific ribosomal subunits and oxidative phosphorylation during embryonic endocrine cell development have not been studied, and thus a fuller interpretation of the importance of these genes remains to be addressed in future studies.

Our scRNA-Seq analysis further showed that relatively few genes that are normally induced during the Preβ to β cell transition are suppressed in *Glis3*KO mice. The expression of *Ins2* is the gene most impacted by the loss of GLIS3 function during this transition indicating that GLIS3 is required for the induction of *Ins2* at this stage. Previous studies have proposed that GLIS3 acts as a pioneer factor for the insulin promoter, with mutations in a GLISBS greatly reducing its transcriptional activation^18,41^. Our snATAC-seq analysis indicated that GLIS3 has a modest effect on chromatin accessibility at the *Ins2* locus (Figure 6B, Supplemental Table 14) and does not have noticeable effects on the chromatin accessibility of other regulated genes (Figure 6C and D). While future studies may seek to confirm these findings by examining histone modifications associated with open/closed chromatin, such techniques are not currently feasible in embryonic single cell populations. We therefore propose that GLIS3 does not act as a pioneer factor but instead as a cooperating transcription factor that contributes to increased accessibility at the *Ins2* promoter in coordination with a pioneer factor (Figure 6E) thereby greatly enhancing *Ins2* transcriptional activation^43^. As we reported previously, although GLIS3 is essential for high *Ins2* expression, it regulates *Ins2* transcription in cooperation with several other islet-enriched transcription factors, such as PDX1, MAFA, and NEUROD1^18^.

Utilizing single cell RNA-sequencing, we were able to identify two novel regulatory functions for GLIS3 in the development of the pancreatic endocrine lineage. First, GLIS3 plays a critical role in the generation/differentiation of BiP cells. The reduction in the number of BiP cells consequently leads to a proportionate reduction in all successive endocrine cell types. Second, GLIS3 is critical for the Preβ to β cell transition. Loss of GLIS3 function generates a unique β-like cell population, in which *Ins2* is greatly repressed and many ribosomal and oxidative phosphorylation genes remain elevated. Dysregulation of these two stages together provide a causal mechanism for the development of neonatal diabetes in GLIS3-deficiency. Future studies should seek to further dissect GLIS3’s early roles in development and possibly reshape our knowledge of how to direct endocrine differentiation for use in stem cell derived therapeutic protocols.

## Acknowledgements

We would like to acknowledge Laura Miller at the NIEHS for her assistance in producing the mice, the Epigenomics and DNA Sequencing Core at NIEHS for carrying out scRNA-seq and snATAC-seq, and the Cell Biology Group at NIEHS for thoughtful comments throughout. AMJ is the guarantor of this work and, as such, had full access to all the data in the study and takes responsibility for the integrity of the data and the accuracy of the data analysis. This research was supported by the Intramural Research Program of the NIEHS, NIH Z01-ES-101585. The contributions of the NIH authors were made as part of their official duties as NIH federal employees, are in compliance with agency policy requirements, and are considered Works of the United States Government. However, the findings and conclusions presented in this paper are those of the author(s) and do not necessarily reflect the views of the NIH or the U.S. Department of Health and Human Services.

## Conflict of Interest

The authors state no conflict of interest.

## Notes

### Competing Interest Statement

The authors have declared no competing interest.

## References

1 Kim, Y. S., Nakanishi, G., Lewandoski, M. & Jetten, A. M. GLIS3, a novel member of the GLIS subfamily of Kruppel-like zinc finger proteins with repressor and activation functions. Nucleic Acids Res 31, 5513–5525 (2003). 10.1093/nar/gkg776

2 Kang, H. S. et al. Transcription factor Glis3: a novel critical player in the regulation of pancreatic β-cell development. Mol Cell Biol 29, 6366–6379 (2009).

3 Watanabe, N. et al. A murine model of neonatal diabetes mellitus in Glis3-deficient mice. FEBS Lett 583, 2108–2113 (2009). 10.1016/j.febslet.2009.05.039

4 Jetten, A. M. GLIS1-3 transcription factors: critical roles in the regulation of multiple physiological processes and diseases. Cell Mol Life Sci 75, 3473–3494 (2018). 10.1007/s00018-018-2841-9

5 Senee, V. et al. Mutations in GLIS3 are responsible for a rare syndrome with neonatal diabetes mellitus and congenital hypothyroidism. Nat Genet 38, 682–687 (2006). 10.1038/ng1802

6 Scoville, D. W., Kang, H. S. & Jetten, A. M. Transcription factor GLIS3: Critical roles in thyroid hormone biosynthesis, hypothyroidism, pancreatic beta cells and diabetes. Pharmacol Ther 215, 107632 (2020). 10.1016/j.pharmthera.2020.107632

7 Dimitri, P. et al. Novel GLIS3 mutations demonstrate an extended multisystem phenotype. Eur. J. Endocrinol 164, 437–443 (2011).

8 London, S. et al. Case Report: Neonatal Diabetes Mellitus Caused by a Novel GLIS3 Mutation in Twins. Front Endocrinol (Lausanne) 12, 673755 (2021). 10.3389/fendo.2021.673755

9 Beak, J. Y., Kang, H. S., Kim, Y. S. & Jetten, A. M. Functional analysis of the zinc finger and activation domains of Glis3 and mutant Glis3(NDH1). Nucleic Acids Res 36, 1690–1702 (2008). 10.1093/nar/gkn009

10 Scoville, D. W. & Jetten, A. M. GLIS3: A Critical Transcription Factor in Islet beta-Cell Generation. Cells 10, 3471 (2021). 10.3390/cells10123471

11 Ding, M. et al. Genetic variants of gestational diabetes mellitus: a study of 112 SNPs among 8722 women in two independent populations. Diabetologia 61, 1758–1768 (2018). 10.1007/s00125-018-4637-8

12 Palmer, N. D. et al. A genome-wide association search for type 2 diabetes genes in African Americans. PloS one 7, e29202 (2012). 10.1371/journal.pone.0029202

13 Yue, S. et al. Genome-wide analysis study of gestational diabetes mellitus and related pathogenic factors in a Chinese Han population. BMC Pregnancy Childbirth 23, 856 (2023). 10.1186/s12884-023-06167-3

14 Meulebrouck, S. et al. Pathogenic monoallelic variants in GLIS3 increase type 2 diabetes risk and identify a subgroup of patients sensitive to sulfonylureas. Diabetologia 67, 327–332 (2024). 10.1007/s00125-023-06035-x

15 Duarte, G. C. K., Assmann, T. S., Dieter, C., de Souza, B. M. & Crispim, D. GLIS3 rs7020673 and rs10758593 polymorphisms interact in the susceptibility for type 1 diabetes mellitus. Acta Diabetol 54, 813–821 (2017). 10.1007/s00592-017-1009-7

16 Sabiha, B. et al. Assessment of genetic risk of type 2 diabetes among Pakistanis based on GWAS-implicated loci. Gene 783, 145563 (2021). 10.1016/j.gene.2021.145563

17 Yang, Y., Chang, B. H. & Chan, L. Sustained expression of the transcription factor GLIS3 is required for normal beta cell function in adults. EMBO Mol Med 5, 92–104 (2013). 10.1002/emmm.201201398

18 ZeRuth, G. T., Takeda, Y. & Jetten, A. M. The Kruppel-like protein Gli-similar 3 (Glis3) functions as a key regulator of insulin transcription. Molecular endocrinology (Baltimore, Md.) 27, 1692–1705 (2013). 10.1210/me.2013-1117

19 Scoville, D. W., Lichti-Kaiser, K., Grimm, S. & Jetten, A. GLIS3 binds pancreatic beta cell regulatory regions alongside other islet transcription factors. J Endocrinol 243, 1–14 (2019). 10.1530/JOE-19-0182

20 Yang, Y., Chang, B. H., Samson, S. L., Li, M. V. & Chan, L. The Kruppel-like zinc finger protein Glis3 directly and indirectly activates insulin gene transcription. Nucleic Acids Res 37, 2529–2538 (2009). 10.1093/nar/gkp122

21 Bele, S., Wokasch, A. S. & Gannon, M. Epigenetic modulation of cell fate during pancreas development. Trends Dev Biol 16, 1–27 (2023).

22 Pan, F. C. & Wright, C. Pancreas organogenesis: from bud to plexus to gland. Dev Dyn 240, 530–565 (2011). 10.1002/dvdy.22584

23 Kang, H. S., Takeda, Y., Jeon, K. & Jetten, A. M. The Spatiotemporal Pattern of Glis3 Expression Indicates a Regulatory Function in Bipotent and Endocrine Progenitors during Early Pancreatic Development and in Beta, PP and Ductal Cells. PLoS One 11, e0157138 (2016). 10.1371/journal.pone.0157138

24 Yang, Y. et al. The Krüppel-like zinc finger protein GLIS3 transactivates neurogenin 3 for proper fetal pancreatic islet differentiation in mice. Diabetologia 54, 2595–2605 (2011).

25 Kim, Y. S. et al. Glis3 regulates neurogenin 3 expression in pancreatic beta-cells and interacts with its activator, Hnf6. Mol Cells 34, 193-200 (2012). 10.1007/s10059-012-0109-z

26 Amin, S. et al. Discovery of a drug candidate for GLIS3-associated diabetes. Nat Commun 9, 2681 (2018). 10.1038/s41467-018-04918-x

27 Kang, H. S., Beak, J. Y., Kim, Y. S., Herbert, R. & Jetten, A. M. Glis3 is associated with primary cilia and Wwtr1/TAZ and implicated in polycystic kidney disease. Mol Cell Biol 29, 2556–2569 (2009). 10.1128/MCB.01620-08

28 Zheng, G. X. et al. Massively parallel digital transcriptional profiling of single cells. Nat Commun 8, 14049 (2017). 10.1038/ncomms14049

29 Hao, Y. et al. Integrated analysis of multimodal single-cell data. Cell 184, 3573–3587 e3529 (2021). 10.1016/j.cell.2021.04.048

30 Germain, P. L., Lun, A., Garcia Meixide, C., Macnair, W. & Robinson, M. D. Doublet identification in single-cell sequencing data using scDblFinder. F1000Res 10, 979 (2021). 10.12688/f1000research.73600.2

31 Young, M. D. & Behjati, S. SoupX removes ambient RNA contamination from droplet-based single-cell RNA sequencing data. Gigascience 9 (2020). 10.1093/gigascience/giaa151

32 Satpathy, A. T. et al. Massively parallel single-cell chromatin landscapes of human immune cell development and intratumoral T cell exhaustion. Nat Biotechnol 37, 925–936 (2019). 10.1038/s41587-019-0206-z

33 Stuart, T., Srivastava, A., Madad, S., Lareau, C. A. & Satija, R. Single-cell chromatin state analysis with Signac. Nat Methods 18, 1333–1341 (2021). 10.1038/s41592-021-01282-5

34 Pliner, H. A. et al. Cicero Predicts cis-Regulatory DNA Interactions from Single-Cell Chromatin Accessibility Data. Mol Cell 71, 858–871 e858 (2018). 10.1016/j.molcel.2018.06.044

35 Zhang, Y. et al. Model-based analysis of ChIP-Seq (MACS). Genome Biol 9, R137 (2008). 10.1186/gb-2008-9-9-r137

36 Dimitri, P. The role of GLIS3 in thyroid disease as part of a multisystem disorder. Best Pract Res Clin Endocrinol Metab 31, 175–182 (2017). 10.1016/j.beem.2017.04.007

37 ZeRuth, G. T., Williams, J. G., Cole, Y. C. & Jetten, A. M. HECT E3 Ubiquitin Ligase Itch Functions as a Novel Negative Regulator of Gli-Similar 3 (Glis3) Transcriptional Activity. PLoS One 10, e0131303 (2015). 10.1371/journal.pone.0131303

38 Larsson, L. I. On the development of the islets of Langerhans. Microsc Res Tech 43, 284–291 (1998). 10.1002/(SICI)1097-0029(19981115)43:4<284::AID-JEMT2>3.0.CO;2-0

39 Yu, X. X. et al. Defining multistep cell fate decision pathways during pancreatic development at single-cell resolution. EMBO J 38, e100164 (2019). 10.15252/embj.2018100164

40 Yang, Y., Bush, S. P., Wen, X., Cao, W. & Chan, L. Differential Gene Dosage Effects of Diabetes-Associated Gene GLIS3 in Pancreatic beta Cell Differentiation and Function. Endocrinology 158, 9–20 (2017). 10.1210/en.2016-1541

41 Akerman, I. et al. Neonatal diabetes mutations disrupt a chromatin pioneering function that activates the human insulin gene. Cell Rep 35, 108981 (2021). 10.1016/j.celrep.2021.108981

42 Bearrows, S. C. et al. Chromogranin B regulates early-stage insulin granule trafficking from the Golgi in pancreatic islet beta-cells. J Cell Sci 132 (2019). 10.1242/jcs.231373

43 Balsalobre, A. & Drouin, J. Pioneer factors as master regulators of the epigenome and cell fate. Nat Rev Mol Cell Biol 23, 449–464 (2022). 10.1038/s41580-022-00464-z

44 Gradwohl, G., Dierich, A., LeMeur, M. & Guillemot, F. neurogenin3 is required for the development of the four endocrine cell lineages of the pancreas. Proc Natl Acad Sci U S A 97, 1607–1611 (2000). 10.1073/pnas.97.4.1607

45 Gu, G., Dubauskaite, J. & Melton, D. A. Direct evidence for the pancreatic lineage: NGN3+ cells are islet progenitors and are distinct from duct progenitors. Development 129, 2447–2457 (2002). 10.1242/dev.129.10.2447

46 Nogueira, T. C. et al. GLIS3, a susceptibility gene for type 1 and type 2 diabetes, modulates pancreatic beta cell apoptosis via regulation of a splice variant of the BH3-only protein Bim. PLoS Genet 9, e1003532 (2013). 10.1371/journal.pgen.1003532

47 Dooley, J. et al. Genetic predisposition for beta cell fragility underlies type 1 and type 2 diabetes. Nat Genet 48, 519–527 (2016). 10.1038/ng.3531

48 Xue, S. et al. RNA regulons in Hox 5’ UTRs confer ribosome specificity to gene regulation. Nature 517, 33–38 (2015). 10.1038/nature14010

49 Luan, Y. et al. Deficiency of ribosomal proteins reshapes the transcriptional and translational landscape in human cells. Nucleic Acids Res 50, 6601–6617 (2022). 10.1093/nar/gkac053

50 Jia, X. et al. Protein translation: biological processes and therapeutic strategies for human diseases. Signal Transduct Target Ther 9, 44 (2024). 10.1038/s41392-024-01749-9

