## Supplemental Figures for "Identification of distinct functions of GLIS3 in β-cell generation critical to prevention of neonatal diabetes"

### Slide 1
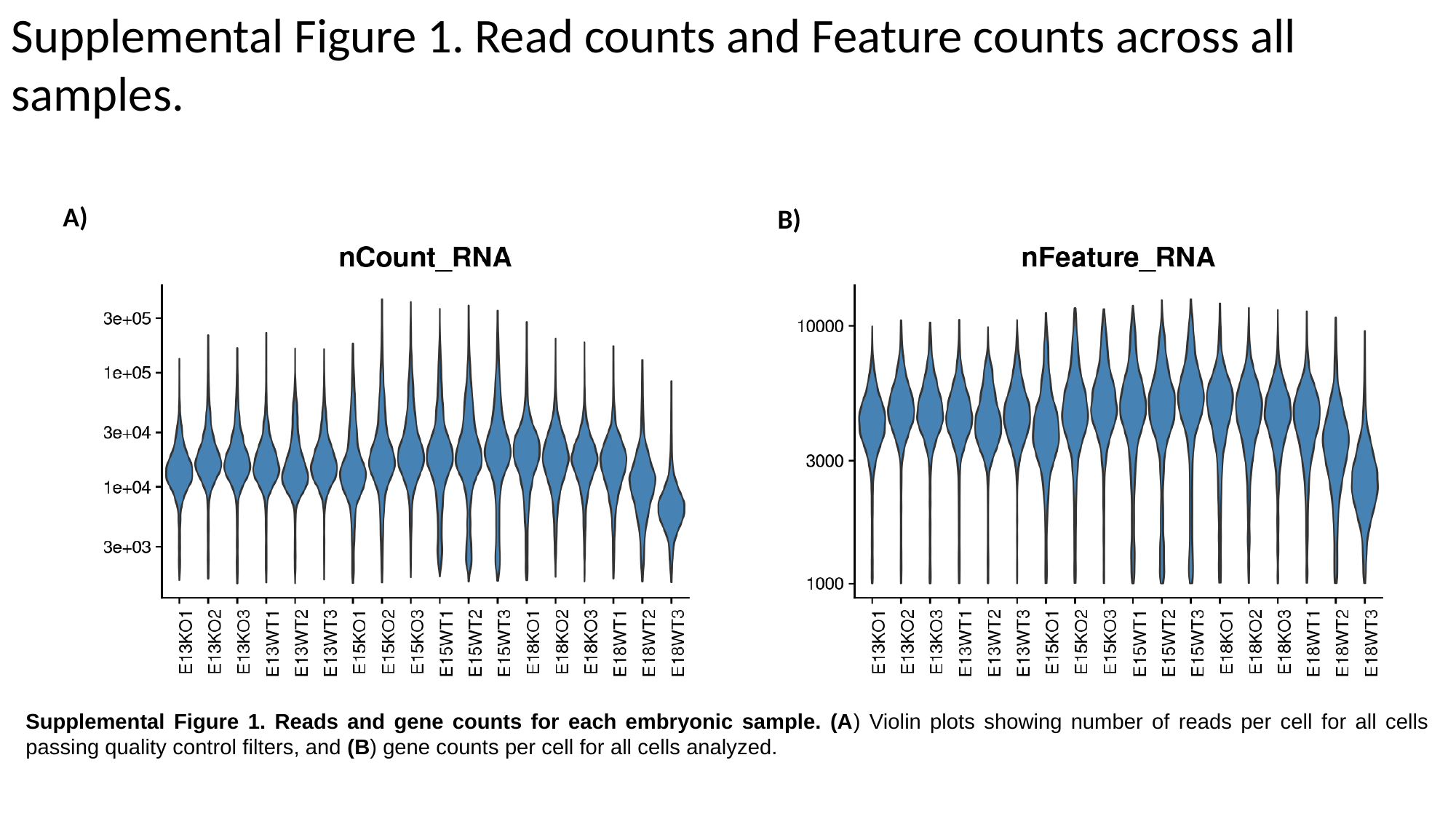

Supplemental Figure 1. Read counts and Feature counts across all samples.
A)
B)
Supplemental Figure 1. Reads and gene counts for each embryonic sample. (A) Violin plots showing number of reads per cell for all cells passing quality control filters, and (B) gene counts per cell for all cells analyzed.

### Slide 2
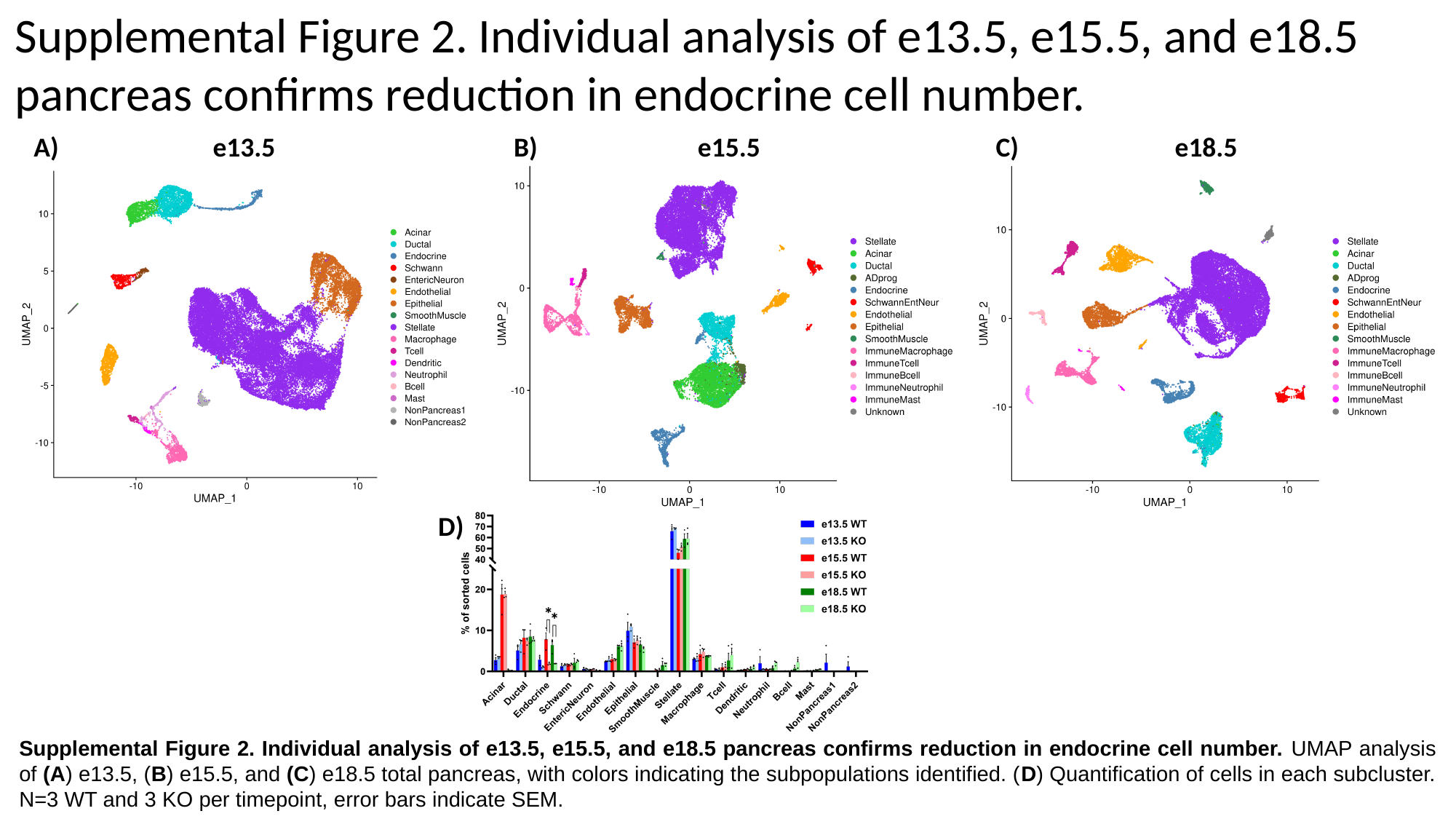

Supplemental Figure 2. Individual analysis of e13.5, e15.5, and e18.5 pancreas confirms reduction in endocrine cell number.
e13.5
A)
e15.5
B)
e18.5
C)
D)
Supplemental Figure 2. Individual analysis of e13.5, e15.5, and e18.5 pancreas confirms reduction in endocrine cell number. UMAP analysis of (A) e13.5, (B) e15.5, and (C) e18.5 total pancreas, with colors indicating the subpopulations identified. (D) Quantification of cells in each subcluster. N=3 WT and 3 KO per timepoint, error bars indicate SEM.

### Slide 3
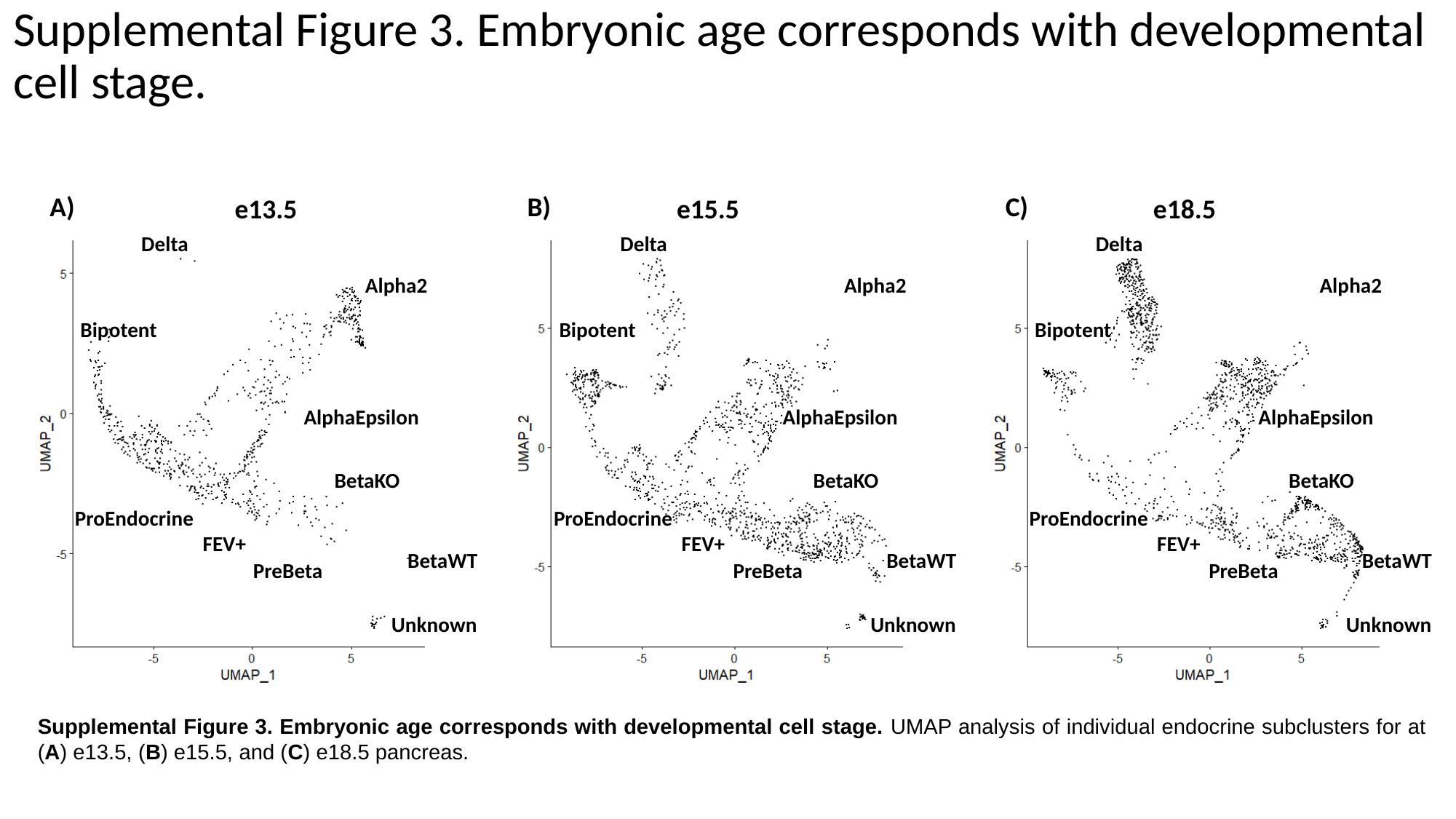

Supplemental Figure 3. Embryonic age corresponds with developmental cell stage.
B)
C)
A)
e13.5
e15.5
e18.5
Delta
Delta
Delta
Alpha2
Alpha2
Alpha2
Bipotent
Bipotent
Bipotent
AlphaEpsilon
AlphaEpsilon
AlphaEpsilon
BetaKO
BetaKO
BetaKO
ProEndocrine
ProEndocrine
ProEndocrine
FEV+
FEV+
FEV+
BetaWT
BetaWT
BetaWT
PreBeta
PreBeta
PreBeta
Unknown
Unknown
Unknown
Supplemental Figure 3. Embryonic age corresponds with developmental cell stage. UMAP analysis of individual endocrine subclusters for at (A) e13.5, (B) e15.5, and (C) e18.5 pancreas.

### Slide 4
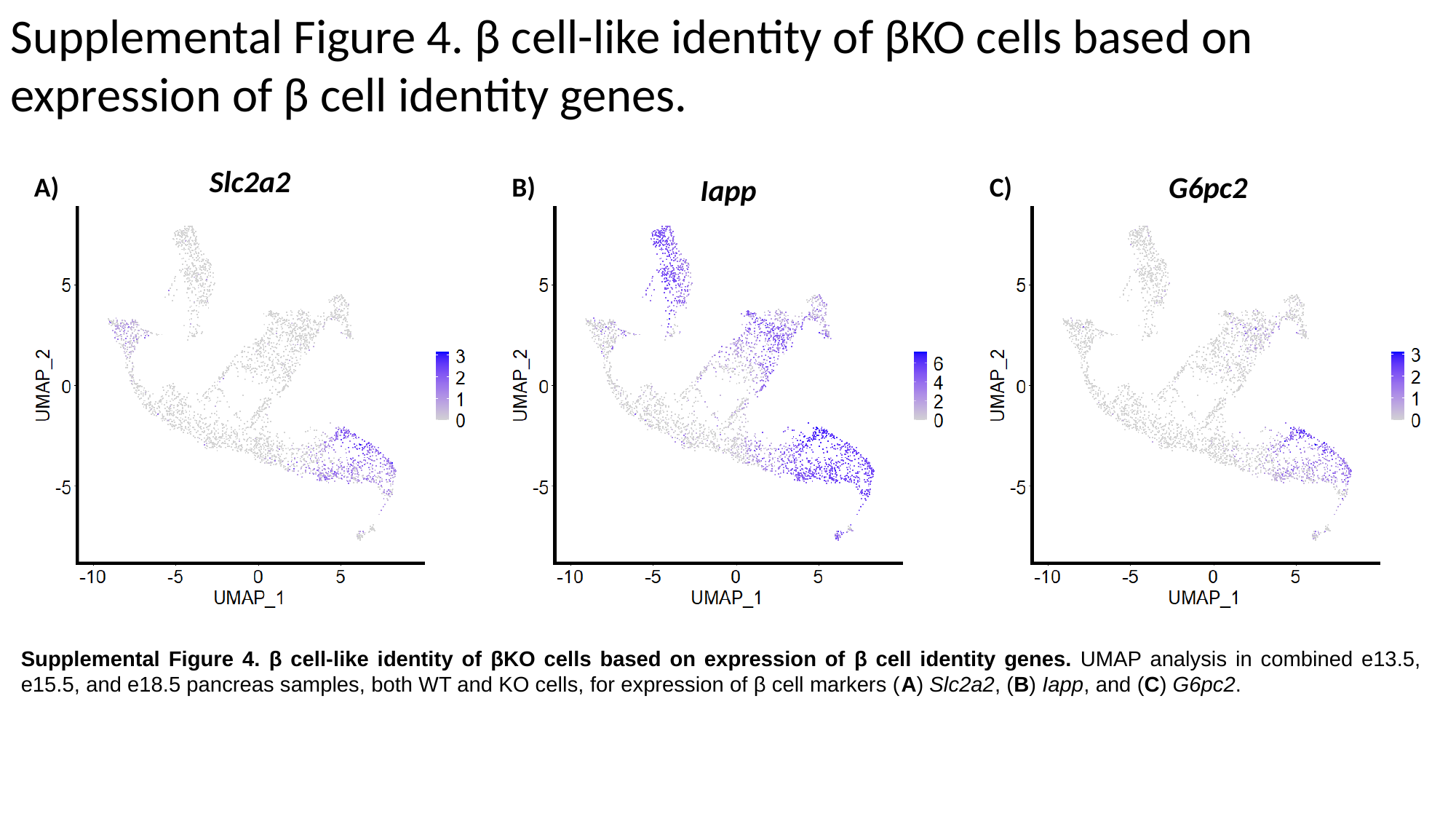

Supplemental Figure 4. β cell-like identity of βKO cells based on expression of β cell identity genes.
Slc2a2
G6pc2
B)
C)
A)
Iapp
Supplemental Figure 4. β cell-like identity of βKO cells based on expression of β cell identity genes. UMAP analysis in combined e13.5, e15.5, and e18.5 pancreas samples, both WT and KO cells, for expression of β cell markers (A) Slc2a2, (B) Iapp, and (C) G6pc2.

### Slide 5
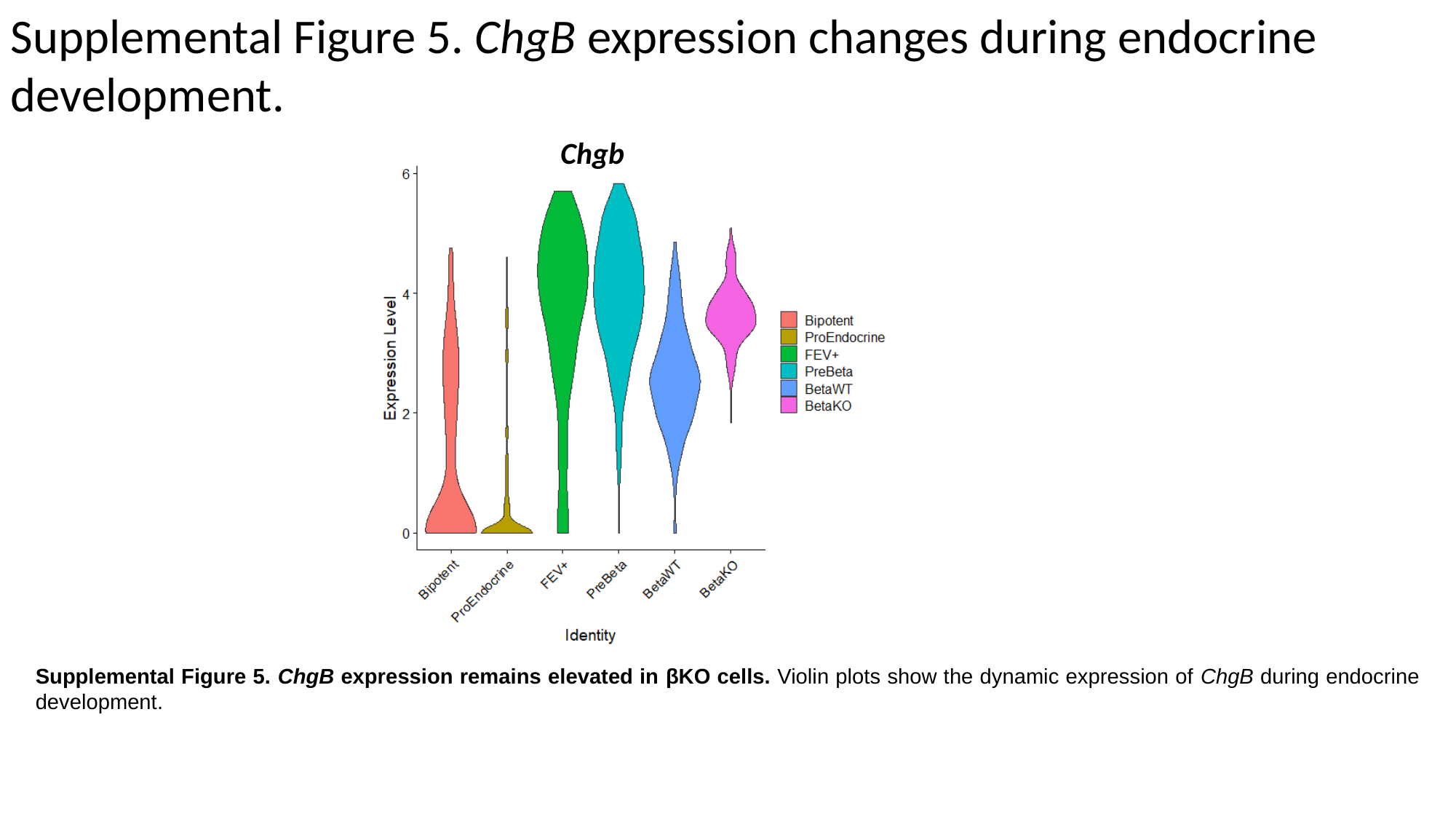

Supplemental Figure 5. ChgB expression changes during endocrine development.
Chgb
Supplemental Figure 5. ChgB expression remains elevated in βKO cells. Violin plots show the dynamic expression of ChgB during endocrine development.

### Slide 6
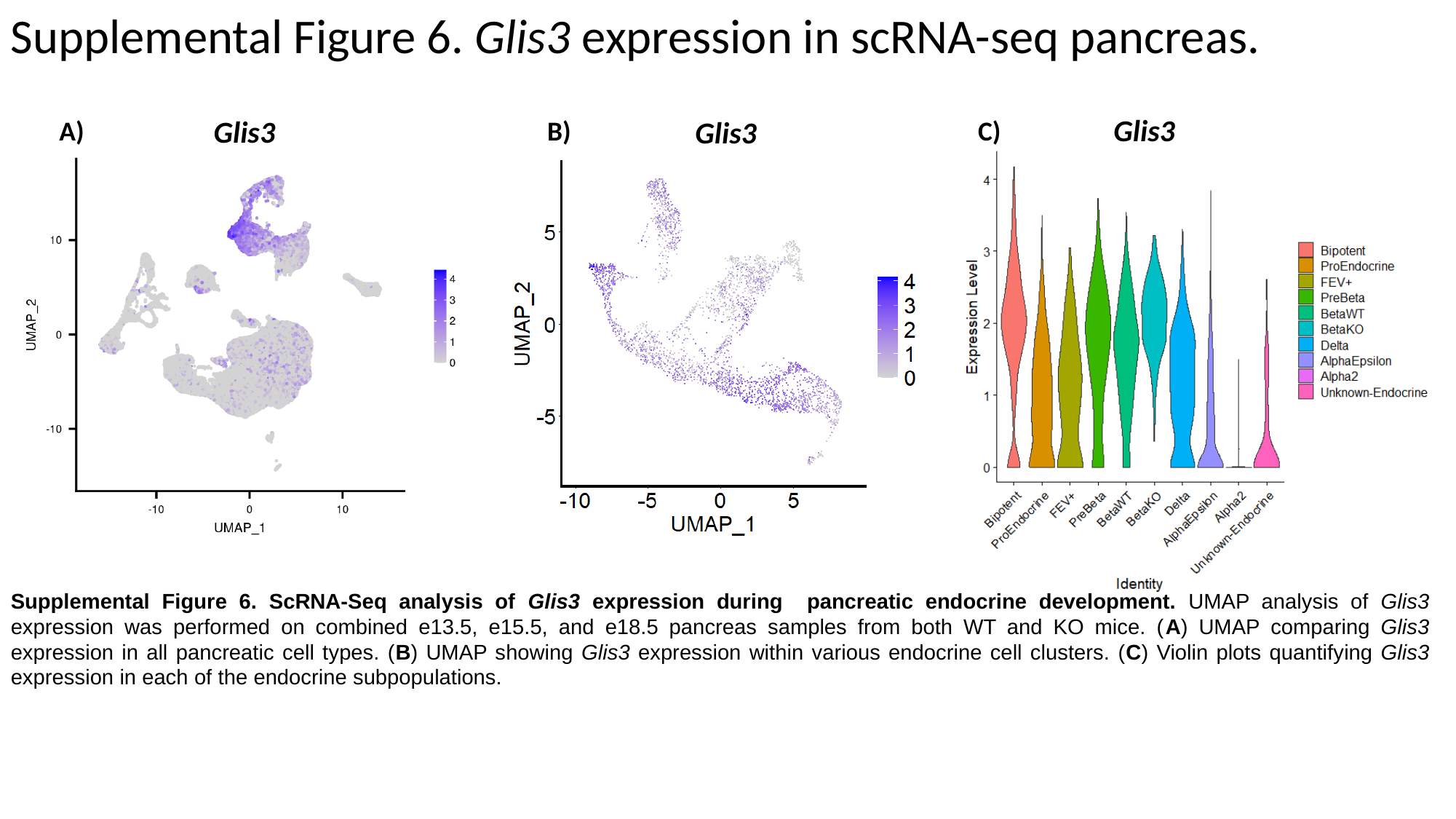

Supplemental Figure 6. Glis3 expression in scRNA-seq pancreas.
Glis3
Glis3
A)
Glis3
B)
C)
Supplemental Figure 6. ScRNA-Seq analysis of Glis3 expression during pancreatic endocrine development. UMAP analysis of Glis3 expression was performed on combined e13.5, e15.5, and e18.5 pancreas samples from both WT and KO mice. (A) UMAP comparing Glis3 expression in all pancreatic cell types. (B) UMAP showing Glis3 expression within various endocrine cell clusters. (C) Violin plots quantifying Glis3 expression in each of the endocrine subpopulations.

### Slide 7
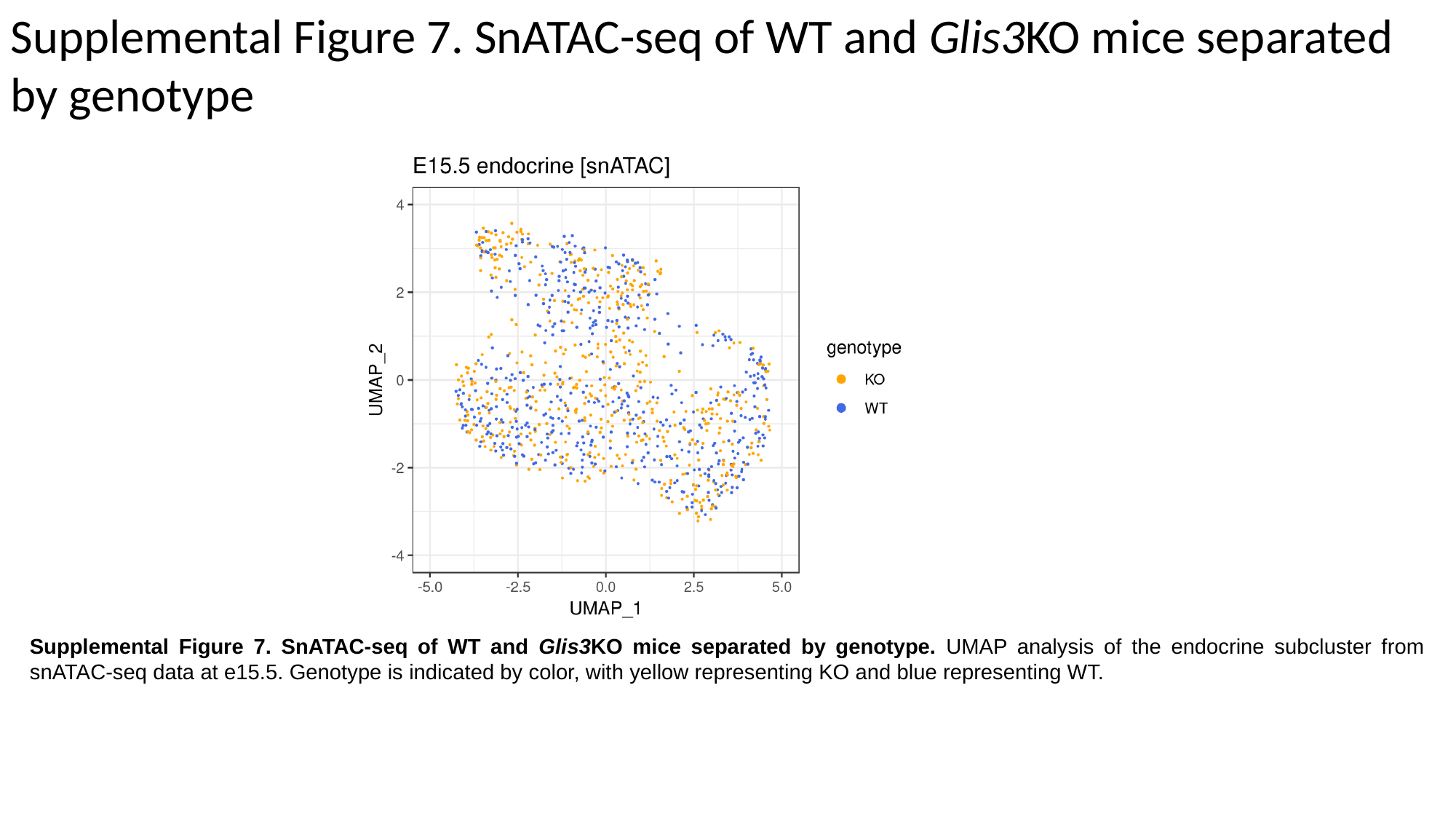

Supplemental Figure 7. SnATAC-seq of WT and Glis3KO mice separated by genotype
Supplemental Figure 7. SnATAC-seq of WT and Glis3KO mice separated by genotype. UMAP analysis of the endocrine subcluster from snATAC-seq data at e15.5. Genotype is indicated by color, with yellow representing KO and blue representing WT.
